# Neuromodulatory organization in the developing rat somatosensory cortex

**DOI:** 10.1101/2022.11.11.516108

**Authors:** Cristina Colangelo, Alberto Muñoz, Alberto Antonietti, Alejandro Antón-Fernández, Armando Romani, Joni Herttuainen, Henry Markram, Javier DeFelipe, Srikanth Ramaswamy

**Author notes:** Correspondence: Srikanth Ramaswamy. Cristina Colangelo and Alberto Muñoz contributed equally.

## Abstract

The vast majority of cortical synapses are found in the neuropil which is implicated in multiple and diverse functions underlying brain computation. Unraveling the organizing principles of the cortical neuropil requires an intricate characterization of synaptic connections established by excitatory and inhibitory axon terminals, of intrinsic and extrinsic origin and from ascending projections that govern the function of cortical microcircuits through the release of neuromodulators either through point-to-point chemical synapses or diffuse volume transmission (VT). Even though neuromodulatory release has been studied for almost a century it is still not clear if one modality prevails upon the other. The hindlimb representation of the somatosensory cortex (HLS1) of two-week old Wistar rats has served as a model system to dissect the microcircuitry of neurons and their synaptic connections. In the present study, we quantified the fiber length per cortical volume and the density of varicosities for cholinergic, catecholaminergic and serotonergic neuromodulatory systems in the cortical neuropil using immunocytochemical staining and stereological techniques. Acquired data were integrated into a novel computational framework to reconcile the specific modalities and predict the effects of neuromodulatory release in shaping neocortical network activity. We predict that acetylcholine (ACh), dopamine (DA), serotonin (5-HT) release desynchronizes cortical activity by inhibiting slow oscillations (delta range), and that 5-HT triggers faster oscillations (theta). Moreover, we found that high levels (>40%) of neuromodulatory VT are sufficient to induce network desynchronization, but also that combining volume release with synaptic inputs leads to more robust and stable effects, meaning that lower levels of VT are needed to achieve the same outcome (10%).

## Introduction

Knowledge of the principal organization of the neocortex requires the characterization of the finest details of the complexity of the cortical neuropil. This includes the characterization of the density and distribution of the cortical synaptic junctions, most of which (90-98%) are established in the neuropil (DeFelipe et al., 1999). Ascending cholinergic, catecholaminergic and serotoninergic regulatory systems made by intricate networks of varicose fibers participate in the control of cortical microcircuits both through synaptic contacts (“classical” point-to-point chemical synapses), and non-synaptic diffuse or volume transmission (VT) processes, impacting cortical states, functions and development (Mechawar et al., 2002; Fuxe et al., 2010; Rho et al., 2018). The hindlimb representation of the somatosensory cortex (HLS1) of the two-week old Wistar rats represents an excellent model to characterize the organizational principles of neocortical microcircuits due to its accessibility and the availability of detailed anatomical, molecular and physiological experimental data on its cellular and synaptic organization. In the last few years it has received much attention within the framework of the Blue Brain Project (http://bluebrain.epfl.ch; https://cajalbbp.es) (Markram et al., 2015). Modeling the influence that ascending modulatory systems exert on cortical microcircuitry requires a detailed assessment of the density and distribution of cholinergic, catecholaminergic and serotoninergic fibers and varicosities compared to all types of synapses. In the present study we have estimated the densities and laminar distribution patterns of these modulatory systems using immunostaining and stereological techniques. Furthermore, to extend our characterization of the neuromodulatory innervation of the developing rodent neocortex, we use computational methods to digitally reconstruct cholinergic, dopaminergic and serotoninergic virtual fibers in a detailed, biologically accurate model of the cortical column (Markram et al., 2015). Our algorithmic approach to reconstruct neuromodulatory innervation leverages experimental datasets to predict missing biological information such as the relative proportions of targeted neurons or the number of contacts established by each neuromodulatory fiber. We subsequently simulate the activation of acetylcholine (ACh), dopamine (DA), serotonin (5-HT), projections to model the synaptic and volumetric release of neuromodulators in the neocortical sensory microcircuit, and predict a desynchronizing effect on the network. We found that synaptic and volumetric release of neuromodulators have synergistic effects and lead to more robust desynchronizing effects when combined, rather than on their own. Moreover, we predict that ACh and DA inhibit slow oscillations, whereas 5-HT additionally brings about faster oscillatory activity. Overall, we propose a framework to integrate what is known about synaptic and non-synaptic transmission to attempt to reconcile conflicting reports in the literature and direct experimental research to answering long-standing questions about the nature of neuromodulatory release.

## Material and Methods

### Animals

Wistar rats (n=11, aged 14 days) were sacrificed by administering a lethal intraperitoneal injection of sodium pentobarbital (40 mg/kg), and they were then perfused intracardially with saline solution followed by 4% paraformaldehyde in 0.1 M phosphate buffer (PB), pH 7.4. All experiments were approved by the local ethics committee of the Spanish National Research Council (CSIC) and performed in accordance with the guidelines established by the European Union regarding the use and care of laboratory animals (Directive 2010/63/EU). Brains were removed and post-fixed by immersion in the same fixative for 7h at 4ºC. For the quantification of the cholinergic, catecholaminergic and serotoninergic fibers, after post-fixation, six brains (Rabb6-Rabb11) were cryoprotected in 30% sucrose solution in PB until they sank, frozen in dry ice and cut in the coronal plane with a sliding freezing microtome. In all cases, 50 μm-thick sections extending in the antero-posterior axis from the anterior commissure to the rostral limit of the hippocampal formation and including the hindlimb representation area of the primary somatosensory cortex (Paxinos and Watson, 2007) were processed for immunocytochemistry.

### Immunocytochemistry

The sections were rinsed in PB and to block non-specific antibody binding they were preincubated for 1 h at room temperature in a stock solution containing 3% normal serum of the species in which the secondary antibodies were raised (Vector Laboratories, Burlingame, CA) in PB with Triton X-100 (0.25%). After preincubation, sections were incubated for 48 h at 4ºC in the same stock solution containing rabbit anti-serotonin (1:1000, Diasorin, Italy), mouse-anti-Tyrosine hydroxylase (TH, 1:1000, Diasorin) or goat-anti choline acetyltransferase (Chat, 1: 1000, Santa Cruz CA, USA). Sections were then rinsed in PB and incubated in anti-rabbit, anti-mouse, or anti-goat biotinylated secondary antibodies (1:200; Vector Laboratories, Burlingame, CA). After rinsing in PB, a first set of sections of each type of immunostaining was processed for immunofluorescence being incubated for 2 h at room temperature in Alexa 488-coupled Streptavidin (1:200; Molecular Probes, Eugene, OR, USA). Sections were rinsed and stained with DAPI to reveal borders between layers and cytoarchitectonic areas. The sections were then washed in PB, mounted in antifade mounting medium (Invitrogen/Molecular Probes, Eugene, OR) and studied by conventional fluorescence and confocal microscopy (Zeiss, 710). For confocal microscopy, stripes through the layers of the hindlimb primary somatosensory cortex were scanned from every animal. Z sections were recorded at 0.45 μm intervals through separate channels using a 40x oil-immersion lens (numerical aperture of 1.3). Subsequently, ZEN software (Zeiss) was used to construct composite images from each optical series by combining the images recorded through the different channels. Adobe Photoshop CS4 software was used to generate the figures (Adobe Systems Inc., San Jose, CA). For DAB immunostaining, a second set of sections was processed using the Vectastain ABC immunoperoxidase kit (Vector). Antibody labeling was visualized with 0.05% 3,39-diaminobenzidine tetrahydrochloride (Sigma, St Louis, MO) and 0.01% hydrogen peroxide. The sections were rinsed in PB and mounted on siliconized glass slides. After attachment, sections were lightly Nissl-stained with thionin, dehydrated, cleared with xylene, and the cover slipped. Controls were included in all the immunocytochemical procedures, either by replacing the primary antibodies with preimmune goat serum in some sections, by omitting the secondary antibodies, or by replacing the secondary antibody with an inappropriate secondary antibody. No significant immunolabeling was detected under these control conditions.

### Estimation of fiber length

The fiber length and density of varicosities of the cholinergic, catecholaminergic, and serotonergic systems, DAB-immunostained through the depth of the tissue, were stereologically estimated in every cortical layer using respectively the space ball probe (dissector height of 11-18 μm) and the optical fractionator tool (dissector height of 11 μm in all cases) of Stereo Investigator software (StereoInvestigator 7.0, MicroBrightField Inc. Vermont, USA) following previous studies (Mouton et al., 2002). An oil immersion x100 objective (numerical aperture of 1.35) on a BX51 Olympus microscope equipped with a Prior motorized stage and a JVC video camera was used. The light Nissl staining of every immunostained section helped to distinguish areal and layer limits and to trace the contour lines corresponding to the individual cortical layers within the hindlimb cortex with the aid of an x20 objective. For each cortical layer, type of immunostaining and animal, the number of sampling sites performed and the number sections used was determined by the constraints of maintaining the coefficient of error below 0.09 (Gundersen, 1988). The fiber length and density of varicosities were corrected for shrinkage as brain tissue shrinks during processing. To estimate the shrinkage in our samples we measured the surface area and thickness of the sections using Adobe Photoshop and Stereo Investigator software respectively before and after tissue processing either for immunoperoxidase or immunofluorescence. The surface area after processing was divided by the value before processing to obtain an area shrinkage factor (p2) of 0.89. In addition, the obtained linear shrinkage factor in the z-axis (pZ) was 0.28. Therefore, the volume shrinkage factor (p3=p2·pZ) was 0.25.

### Synaptic model of neuromodulatory release

To build a model of neuromodulatory release we used an in-house tool described in (Markram et al., 2015) that was developed to model projections to the neocortical microcircuit in a way that satisfies experimental constraints. Data about the density of synaptic boutons, or, more specifically in our case, neuromodulatory varicosities is used to constrain the generation of new synaptic release sites in the model, that is, we instantiate a layer-wise varicosity density profile in the model, that matches the experimentally measured density. The neuromodulatory projections approach happens in three major steps: sample, assignment to fiber and parameter selection.

#### Sampling

all morphological segments contained in every layer are pooled; from this distribution, random segments are repeatedly sampled and new synaptic release sites (sRS) are placed at their centers in order to match the experimental constraints. Drawing is performed with replacement (i.e., a segment could be drawn more than once). The probability of drawing a given segment is proportional to its length (where longer segments would be drawn more often).

#### Assignment to fiber

a set of ‘fibers’ is given as input to the whole process. These are a set of vectors with starting positions and directions. The tool creates straight synthetic fibers: in the circuit used for this study these begin in L6 and point straight up through all the layers. The parameter ‘number of fibers’ can be estimated or directly constrained from experimental data. The distances between the newly instantiated sRS and the fibers are used to compute a Gaussian that determines the probability of being picked, and using this probability, a fiber is associated to each segment, or ‘mapped’. Given that we do not directly model the subcortical nuclei from which the projections originate, the fibers are activated via an artificially injected spike train of the desired frequency and duration.

#### Parameter selection

in this step, we select the parameters that are better suited to mimic neuromodulatory connections. Our model of synaptic connections follows a conductance-based approach and comprises several parameters. We redirect the reader to (Markram et al., 2015), and to https://bbp.epfl.ch/nmc-portal/microcircuit.html for an exhaustive description. In the following paragraph we will focus on describing how we constrained the connection parameters from literature reported data.

We reasoned that since the reported number of cholinergic neurons in the rat nucleus basalis of Meynert (NBM) is 7312 (Miettinen et al., 2002) and the NBM projects mainly to S1 (Chaves-Coira et al., 2016, 2018) then there would be 7132 / 26 = 283 fibers projecting to our reconstructed microcircuit, because the entire S1 area is 26 times bigger. Similarly, we assigned 2651 / 26 = 102 fibers to the dopaminergic system, based on cell-counts obtained in the rat ventral tegmental area (VTA) (Nair-Roberts et al., 2008) and estimations of the number of VTA neurons that project to S1 (Aransay et al., 2015). Lastly, we computed the number of serotoninergic fibers; 11500 serotoninergic cell bodies have been counted in the dorsal raphe (DR) according to Descarries and others, and only ∼12% project to the S1 region (Descarries et al., 1982; Wilson & Molliver, 1991). That is 1380 / 26 = 53 5-HT fibers for the O1 circuit. The values obtained seem to fall within reasonable ranges, considering the estimated density of each neuromodulatory system, but nevertheless they rest on assumptions that have not been proven. We refer the reader to the Discussion section where we list the most important assumptions. Parameters for the synaptic dynamics of the new neuromodulatory connections were drawn from experimental distributions representative of the synaptic types (s-types) discussed in (Markram et al., 2015) (**Table 6)**, except for the offset decay time constants (DTC). The DTCs of NM connections were constrained with the aid of literature reported values: we refer the reader to **Table 6**.

### Volumetric model of neuromodulatory release

Our model of volumetric neuromodulatory transmission relies upon the implementation of the synaptic model explained above. To model VT, we first take the experimentally recorded varicosity density profile and we instantiate new release sites (RS) in order to match this density. Then, every RS is taken as the center of a sphere of radius Rmax, which is the sphere of influence of volume transmission (thus the RS becomes a volumetric RS, or vRS). We chose Rmax = 5 μm for ACh (**Figure 4**) as was computed by Borden and colleagues, because it was recorded in rodent neocortical brain areas, as opposed to other values obtained in the mEC or the retina, which nevertheless are in a similar range (Borden et al., 2020; Jing et al., 2018; Sethuramanujam et al., 2021). Dopaminergic transients have also been measured by means of a synthetic catecholamine nanosensor which revealed DA hotspots with a median size of 2 μm (Beyene et al., 2019) so we instantiated Rmax = 2 μm for DA connections, even though this data was recorded in the striatal region. Serotonin’s volumetric influence was estimated by (Bunin & Wightman, 1998) who recorded serotoninergic extrasynaptic transmission via carbon fiber microelectrodes in the rat DR; thus, we selected Rmax = 3 μm for 5-HT signals. Subsequently all morphological segments within the sphere are sampled and a new conductance is instantiated in a random position along each segment. Here we assume that extrasynaptic cholinergic receptors are equally distributed across neuronal compartments.In this case the vRS is the source of the NM signal, and the segment where the new conductance is placed is the target. Still, these two do not coincide like in the synaptic implementation of NM release. Thus, we developed a way to mimic the one-source-to-many-targets characteristic of VT. The new volumetric connections are parametrized in the same way as the synaptic ones (see above), but the conductance values in this case are scaled according to the distance from the vRS. Specifically, we determine a scaling factor whose value ranges from 1 to 0.1 and is inversely proportional to the distance from the vRS. Here, we assume that the strength of the volumetric signal decreases linearly with the distance from the release site in order to have a simple model of ACh diffusion and catalysis. We refer the reader again to **Table 6** for a more specific description of the DTC values used to parametrize the kinetics of VT.

### Microcircuit

Our *in silico* neuromodulation model was implemented in a digital microcircuit extracted from a large-scale model covering the entirety of the non-barrel primary somatosensory cortex of the juvenile rat (Reimann et al., 2022). The microcircuit is 26 times smaller than the whole S1 region and comprises 163528 cells arranged in 7 cortical columns stacked horizontally.

### Network simulations

Simulations were conducted using proprietary software based on the NEURON simulation environment. Data were output in the form of binary files containing spike times sampled every 0.1 ms for each neuron in the network. Extracellular calcium and potassium concentrations were modeled by considering their phenomenological effects on neurotransmitter release probability and somatic depolarization, respectively. These values were adjusted to mimic an in vivo-like network state, corresponding (empirically) to extracellular calcium and potassium concentrations of 1.7 mM, and 5.0 mM (∼ 95% somatic firing threshold), respectively. Simulations of ascending neuromodulatory inputs were performed near the transition from the synchronous to the asynchronous state, in order to set the initial state of the network to a substantially synchronized activity pattern. In the control case our microcircuit is oscillating at ∼ 2 Hz, a condition that we use to approximate an inactivated brain state. Every simulation was repeated 30 times with different seeds (tied to random number generators that we use in our simulation pipelines) to reproduce trial variability, and a power density analysis was performed to evaluate the frequency contents of the oscillatory activity of our simulated network of neurons. The stimulus to the neuromodulatory projections was delivered as single-pulse or train stimulation of frequencies ranging from 5 to 30 Hz. To recruit different proportions of ascending inputs we selected varying percentages (from 10 to 100%) of the ‘virtual’ afferent fibers. The stimulus was delivered at t = 2000 ms; the single pulse stimulus had a frequency of 150 Hz and a duration of 10 ms and the train stimulus was applied for 2000 ms.

### Supercomputing

A 2-rack Intel supercomputer using dual socket, 2.3 GHz, 18 core Xeon SkyLake 6140 CPUs, with a total of 120 nodes, 348 GB of memory, and 46 TB of DRAM was used to run the simulations and carry out analysis.

### Simulation outputs analysis

All code for analysis was written in Python 3.6. To estimate the power of the signal at different frequencies, we performed a power spectrum density analysis (PSD) by calculating firing rate frequencies and subsequently applying the Welch transformation. The estimation was performed on the first 1000 ms time interval after the stimulus delivery. To compute the delta power we specifically selected the 2 Hz frequencies.

### Statistical analysis

To estimate possible differences in the length and density values obtained in the different cortical layers, a non-parametric test was performed and correction for multiple comparisons was applied. First, we used Friedman’s test and we found that length and varicosity densities differed across layers; post-hoc analysis (Conover’s test) was later performed to compare all layers for the three neuromodulatory systems. Bonferroni’s correction was applied to account for multiple comparisons. To assess the statistical significance of the PSD analysis results, we used a paired T-student test.

## Results

### Cholinergic fibers

In addition to sparsely distributed ChAT-immunoreactive (IR) neuronal cell bodies distributed from layers 2 to 6, ChAT immunostaining revealed the presence of an intricate network of varicose fibers across all cortical layers of the rat P14 HLS1 (**Fig. 1** and **Table 1**). Unfortunately, at a distance from the cell body, ChAT-ir positive processes emanating from the cell body of cortical cholinergic neurons could not be distinguished from the surrounding positive elements, most of which are known to originate in the basal forebrain (Mechawar et al., 2000). Fibers throughout cortical layers were apparently oriented in all directions but in layers 1 and 6 fibers with an orientation parallel to the pial surface were frequently found. The density of cholinergic fibers was relatively homogeneous throughout cortical thickness and laminar differences were statistically significant only between layers 1 and 4 (P-value = 0.048) (**Fig. 1**). Fiber varicosities were also homogeneously found with no significant differences between the different cortical layers (**Fig. 1**).

**Table 1.** Fiber length and density of varicosities. Fiber length and density of varicosities of serotonergic, catecholaminergic, and cholinergic regulatory systems in HLS1 at P14. Measured data were corrected for shrinkage and refer to the total volume of cortical layers.

| Layer | 5-HT-ir fibers |  | TH-ir fibers |  | ChAT-ir fibers |  |
| --- | --- | --- | --- | --- | --- | --- |
|  | Length | Varicosities | Length | Varicosities | Length | Varicosities |
| Mean ± SD | $\mu\text{m} / 1000 \mu\text{m}^3$ | Var / 1000 $\mu\text{m}^3$ | $\mu\text{m} / 1000 \mu\text{m}^3$ | Var / 1000 $\mu\text{m}^3$ | $\mu\text{m} / 1000 \mu\text{m}^3$ | Var / 1000 $\mu\text{m}^3$ |
| I | $0.74 \pm 0.17$ | $0.29 \pm 0.10$ | $5.36 \pm 1.08$ | $0.42 \pm 0.19$ | $7.30 \pm 1.29$ | $0.47 \pm 0.09$ |
| II | $0.56 \pm 0.12$ | $0.22 \pm 0.06$ | $3.89 \pm 0.76$ | $0.24 \pm 0.04$ | $9.48 \pm 2.82$ | $0.38 \pm 0.06$ |
| III | $0.20 \pm 0.04$ | $0.14 \pm 0.009$ | $3.57 \pm 0.7$ | $0.24 \pm 0.06$ | $9.27 \pm 2.55$ | $0.36 \pm 0.06$ |
| IV | $0.13 \pm 0.05$ | $0.11 \pm 0.02$ | $4.92 \pm 0.47$ | $0.28 \pm 0.03$ | $11.72 \pm 1.52$ | $0.37 \pm 0.03$ |
| V | $0.10 \pm 0.04$ | $0.13 \pm 0.039$ | $3.38 \pm 0.56$ | $0.23 \pm 0.02$ | $9.55 \pm 2.72$ | $0.41 \pm 0.11$ |
| VI | $0.13 \pm 0.04$ | $0.12 \pm 0.02$ | $3.20 \pm 0.29$ | $0.21 \pm 0.05$ | $10.63 \pm 2.40$ | $0.32 \pm 0.09$ |

**Figure 1.**
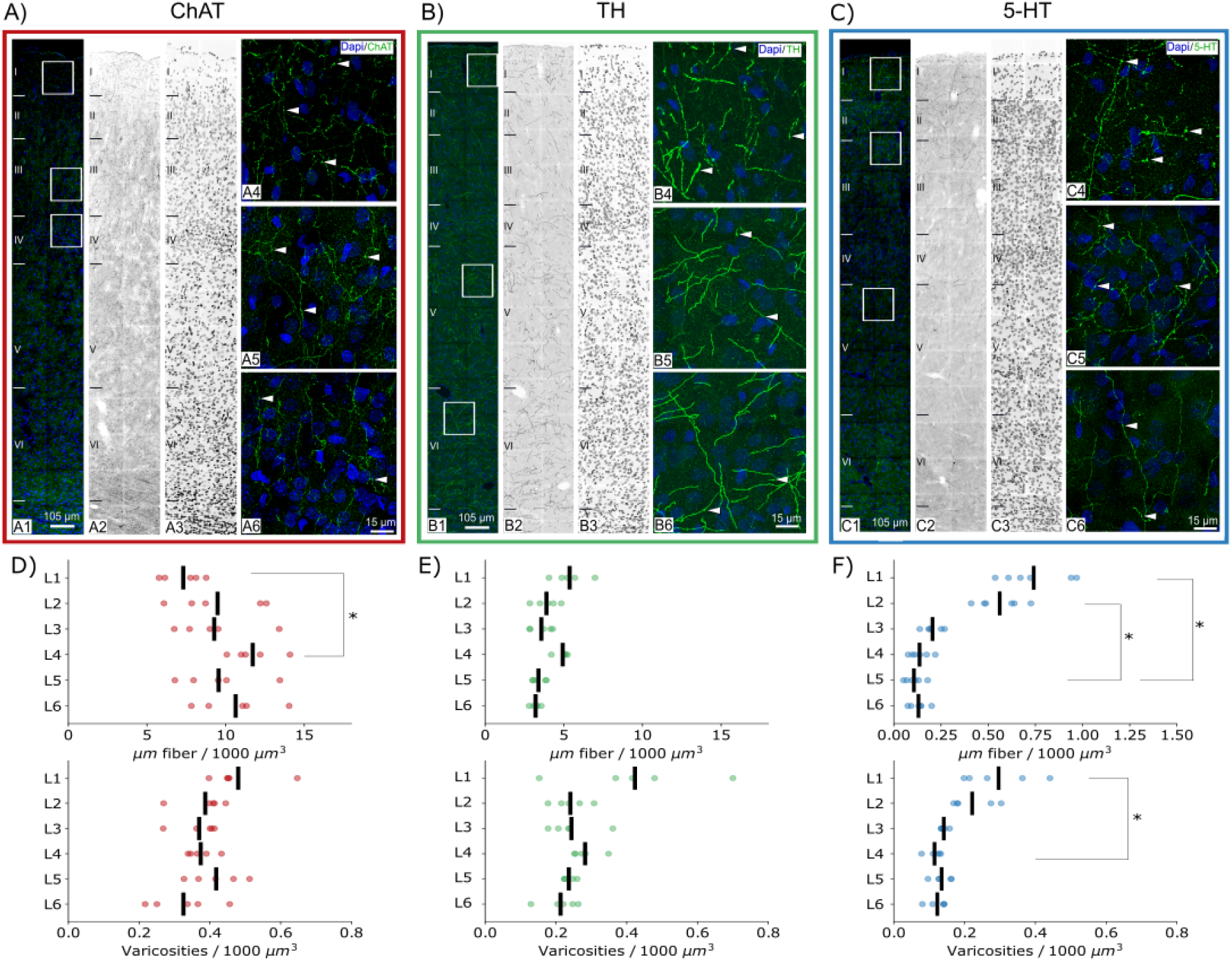
ChAT, 5-HT and TH immunoreactivity. **A)** confocal stack projection image, corresponding to a cortical thickness of 14 μm, showing the distribution of ChAT-immunoreactive fibers (green) in the different layers, as revealed by DAPI staining (blue), of the P14 rat hindlimb somatosensory cortex. A2) and A3) show respectively, in monochrome images, ChAT immunostaining and DAPI staining. Squared zones in A are shown at higher magnification in A4-A5-A6. Arrowheads point to fiber varicosities. **B)** same as in A) but for TH-immunoreactive fibers. **C)** same as in A) but for 5-HT-immunoreactive fibers. **D)** Graph showing the mean values obtained in the HLS1 of the five P14 rats analyzed for the density of fibers (above) and the density of varicosities (below) for the cholinergic system. **E)** same as in D but for the catecholaminergic system. **F)** same as in D but for the serotoninergic system.

**Figure 2.**
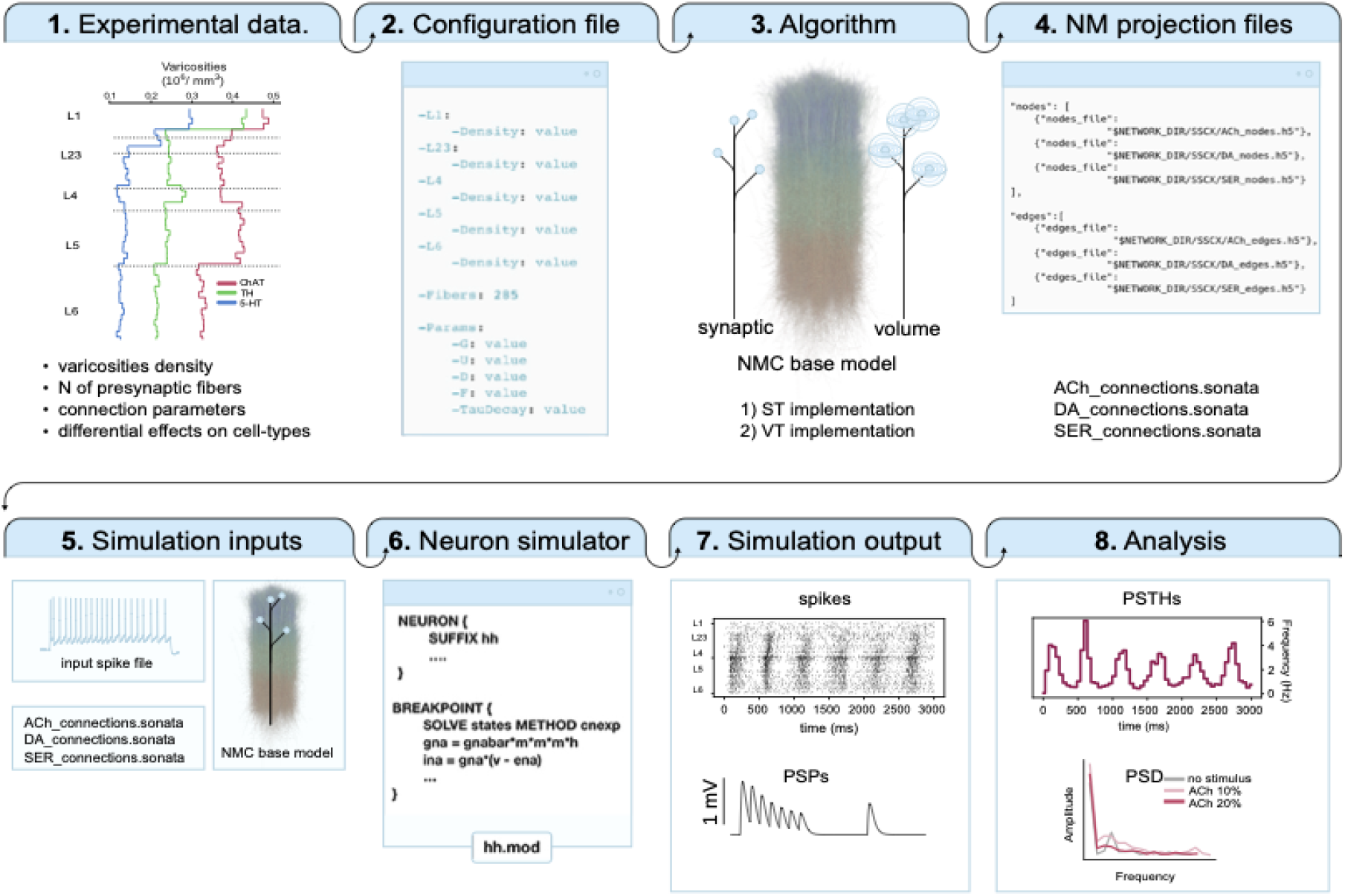
Modeling workflow. ST synaptic transmission; VT volumetric transmission; ACh acetylcholine; DA dopamine, SER serotonin; NMC neocortical microcircuit model; PSP postsynaptic potential; PSTH peristimulus time histogram; PSD power spectral density.

### Catecholaminergic fibers

Immunocytochemistry for tyrosine hydroxylase (TH), the rate-limiting catecholamine synthesizing enzyme, revealed the presence of groups of aspiny non-pyramidal neurons. According to previous studies (Kosaka et al., 1987), these cells are distributed in all cortical layers, although they are most abundant in layers 2–3. The processes arising from this neuronal population might therefore contribute to the TH-ir fibers quantified in the present study (**Fig. 1** and **Table 1**). In addition, TH-ir fibers are considered to label mainly dopaminergic fibers in the cerebral cortex (Lewis et al., 1988; Martin & Spühler, 2013; Sesack et al., 1995), although the possibility that our results include noradrenergic fibers and boutons, mainly labeled with dopamine-β-hydroxylase (Latsari et al., 2002), cannot be excluded. In the rat HLS1 neocortex layers 1 and 4 had the highest density of TH-ir fibers, followed by layers 2 and 3, whereas the density was lowest in layers 5 and 6 (**Fig. 1**). However, no statistical differences were found between layers in the density of TH-ir fibers. The density of TH-ir varicosities followed a similar laminar distribution pattern although no statistical differences were found between layers (**Fig. 1**).

### Serotonergic fibers

Immunocytochemistry for serotonin (5-HT) revealed the presence of numerous 5-HT-ir fibers through all cortical layers (**Fig. 1** and **Table 1**). No immunoreactive cell bodies were found, as expected. A higher density of serotonergic fibers was found in supragranular layers, in layer 1 and 2, compared to layer 5 (p-values respectively 0.007 and 0.035) (**Fig. 1**). Fibers in layers 1 and 6 showed a preferential horizontal orientation parallel to the pial surface whereas in other layers fibers oriented in all spatial orientations but with a radial orientation were frequently found. Fibers with varicosities were also found in all cortical layers and the density of varicosities tended to be higher in superficial rather than in infragranular layers (**Fig. 1**), but a statistically significant difference was found only between L1 and L4 (p-value: 0.048).

### Comparison of the three ascending neuromodulatory systems

The present observations indicate that in terms of fiber length the cholinergic system constitutes the densest neuromodulatory system of the three systems studied, followed by the catecholaminergic fiber system, with the serotonergic system showing the lowest density of fibers. These differences were significant when averaging data from all cortical layers and also in each cortical layer separately (**Fig 1**). Regarding fiber varicosities, that represent the presumed sites of transmitter release, the density of cholinergic varicosities corresponded to 1.4 times the density of varicosities of catecholaminergic varicosities and 2.3 times to the number of serotonin axon varicosities. Finally, the density of catecholaminergic varicosities was 1.6 higher than that of serotonergic varicosities.

### In silico predictions about the organization of neuromodulatory input

We implemented two full sets of neuromodulatory projections that work only with ST and VT respectively. The model allows us to combine the two in the desired proportions, or to only use one projection system at a time. To extend our assessment of the influence of neuromodulatory systems we used the varicosity density profiles across the six neocortical layers to digitally reconstruct cholinergic, dopaminergic and serotoninergic inputs to the hindlimb region of the rat somatosensory cortex and to obtain quantitative anatomical predictions about the columnar targets of the three neuromodulatory innervation systems. Specifically, we estimate the total number of neurons innervated by each projection system, the number of postsynaptic cells contacted by each fiber and the most contacted cell types (**Figure 3**). Overall, we estimated that each cholinergic fiber targets 301.0 ± 493.6 neurons with 336.6 ± 567.4 synaptic RS, while each cortical neuron receives 4.2 ± 2.7 fibers with 4.7 ± 3.3 synaptic RS. In the case of volumetric transmission, we calculated that each ACh fiber contacts up to 7104.4 ± 4581.0 neurons. The most contacted excitatory cell type is the TPC:A in L23, while the most contacted inhibitory cell-type is the LBC in L23. Moreover, we predict that each TH immuno-positive fiber targets 521.4 ± 852.2 neurons with 631.4 ± 1063.4 synaptic RS, while each cortical neuron receives 2.7 ± 1.6 fibers with 3.3 ± 2.3 synaptic RS. In the VT implementation each TH fiber targets 4377.7 ± 3982.9 neurons. The most densely innervated cell-type is the L23 TPC:A while in terms of inhibitory cells the most contacted type is the LBC in L6. Each 5-HT fiber targets 365.7 ± 566.7 neurons with 458.2 ± 740.0 synaptic RS and each neuron receives 2.0 ± 1.1 fibers with 2.5 ± 1.7 synaptic RS. In the VT case serotoninergic contacts reach 3463.2 ± 2605.1 neurons. The most contacted excitatory neuron is the TPC:A in L5 and the most contacted interneuron is the L23 LBC. For a more refined breakdown of the proportions of cell-types innervated by the three systems see panel C in **Figure 3**. Next, we tested whether our virtual neuromodulatory projection systems can be activated to induce network effects reported in the literature. We reasoned that albeit sparse, if properly connected and parameterized, neuromodulatory fibers should elicit a modulation of network activity.

**Figure 3.**
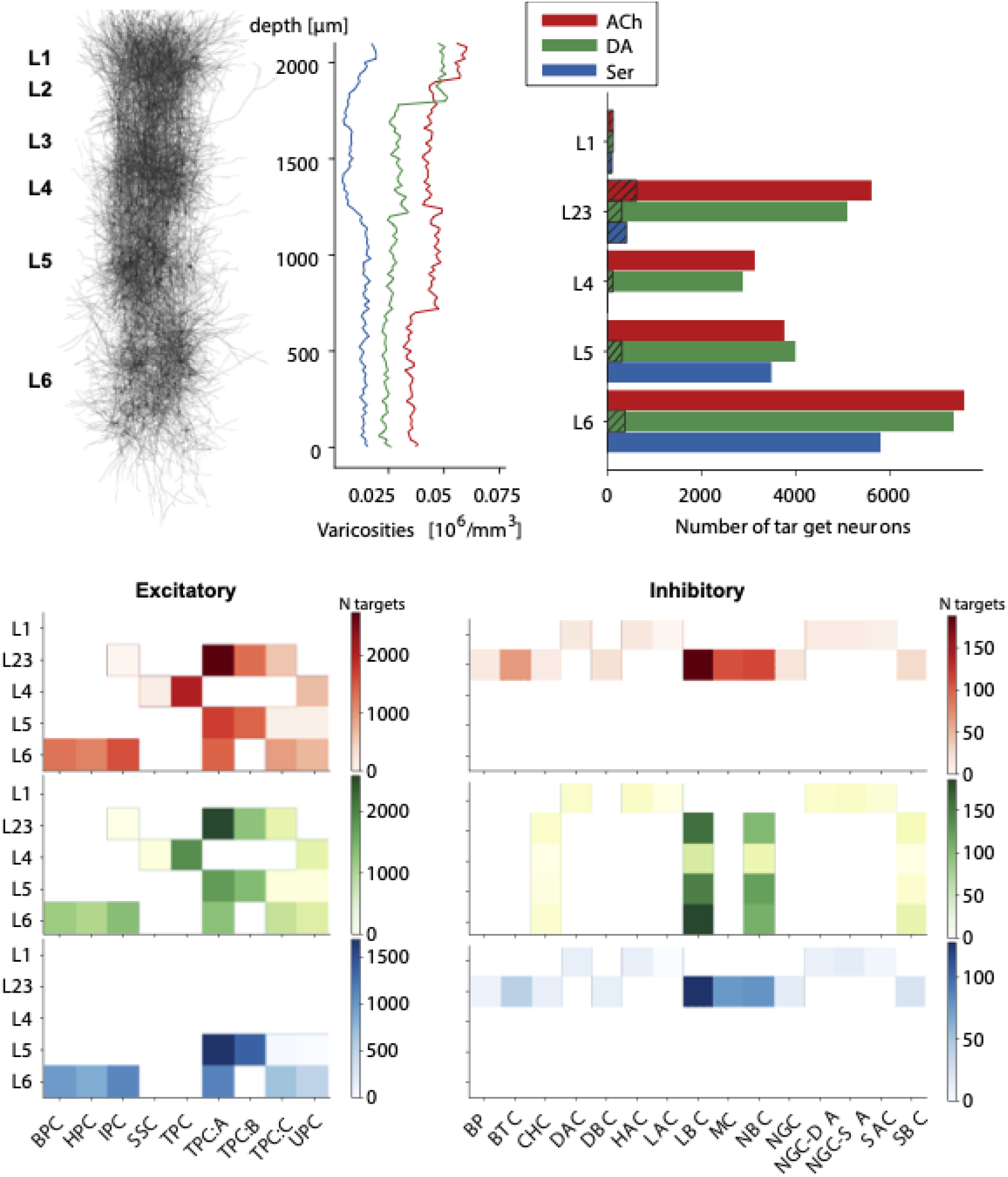
In silico predictions of ChAT, 5-HT and TH. All data is representative of a single column in our in silico neocortical microcircuit. **A)** From left to right: in silico Golgi stain microphotograph of a cortical column; in silico neuromodulatory varicosities densities. **B)** Histograms showing the distribution of the number of postsynaptic targets contacted by each neuromodulatory fiber, for the three neuromodulatory systems. **C)** Heatmap of the proportion of excitatory morphological types contacted by each projection system. **D)** Heatmap of the proportion of inhibitory morphological types contacted by each projection system.

### Simulating cholinergic network effects

Optogenetic activation of ChAT-positive neurons in the basal forebrain (BF) is known to produce a desynchronizing effect on the activity of neurons in sensory cortices (S.-H. Lee & Dan, 2012; Pinto et al., 2013), even though exactly how this is achieved remains unclear. We therefore gathered and integrated data about the effects of optogenetic cholinergic BF neurons stimulation on neocortical cell types residing in sensory cortices, to simulate the activation of cholinergic projections in our detailed model of a neocortical column. As reported in **Table 3** and **Figure 4**, in our implementation, cholinergic inputs depolarize L1 interneurons, L23 distal-targeting interneurons and L5 - L6 pyramidal cells (PCs), and instead have a hyperpolarizing effect on L23 - L4 PCs and L23 proximal-targeting interneurons. The remaining cell-types are not targeted by our ACh connections because cholinergic stimuli in a physiologically relevant range (achieved via optogenetic tools or relatively low concentrations of ACh agonists, i.e., not greater than 100 μm) fail to elicit a response in inhibitory interneurons located in deeper layers (Arroyo et al., 2012; Obermayer et al., 2018). For a more detailed explanation of the parameters used for simulations, we redirect the reader to **Table 6** First, we simulated the all-synaptic activation of the whole cholinergic projection system at increasingly higher stimulation rates (5, 10, 15, 20, 25, 30 Hz) to model progressively higher levels of ACh release. Cholinergic ST inputs significantly reduce the delta component of the power spectrum of network activity (at 20 Hz ST we observe a 49% reduction with respect to control; p-value = 0.18·10^−30^; moreover, this reduction tends to be larger as the stimulation frequency increases. For all subsequent simulations of cholinergic inputs, we therefore chose to mimic the firing frequencies used in optogenetics experiments (around 20 Hz) which in turn imitate the synchronous spiking of cholinergic neurons in the BF (Hedrick & Waters, 2015).

**Figure 4.**
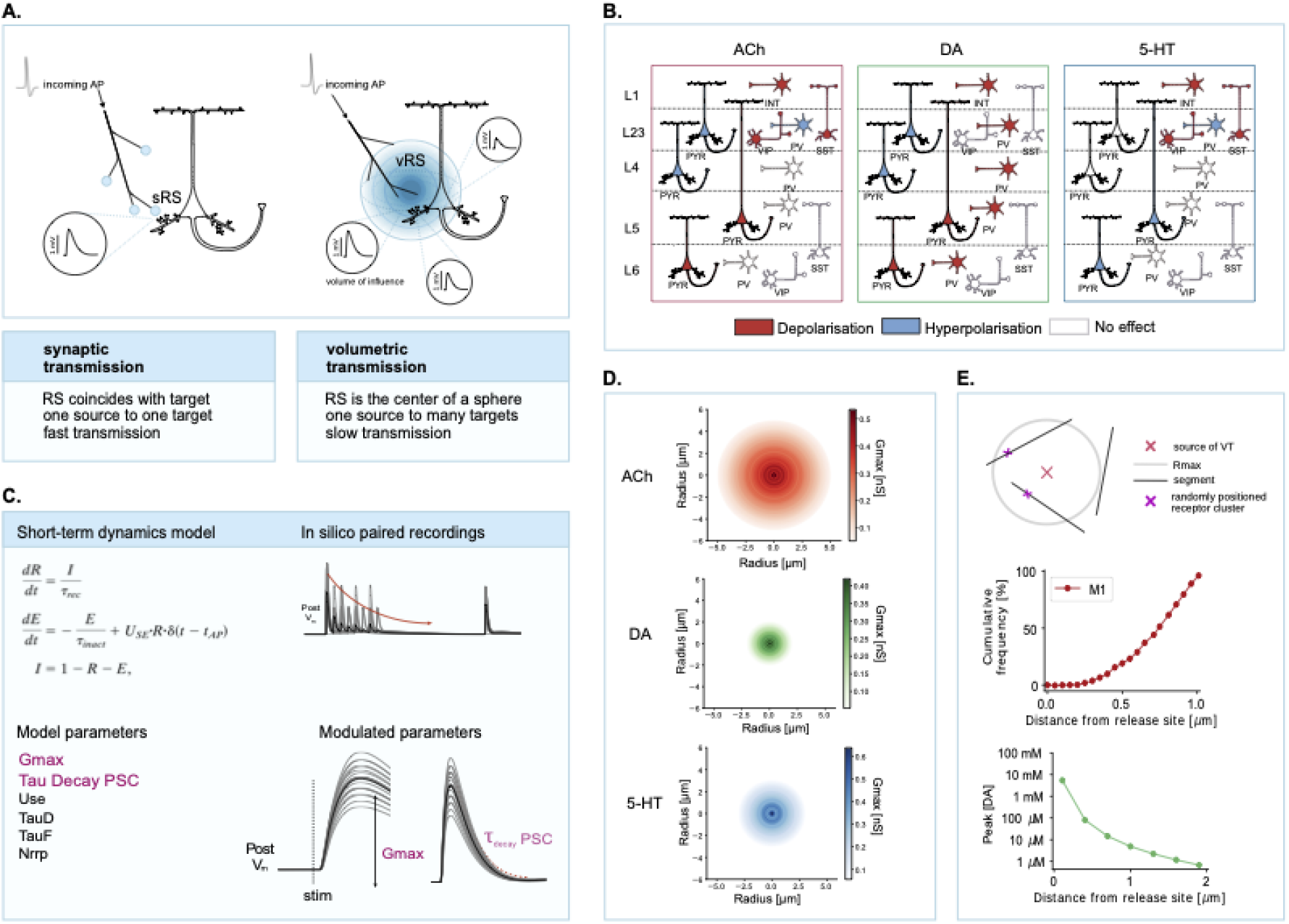
VT vs ST implementation. **A)** Schematics illustrating the differential implementation of volumetric vs synaptic transmission. sRS, synaptic release site, vRS, volumetric release site. **B)** Schematics illustrating the data gathered in **Tables 3-4-5** on the effects of neuromodulator release on a given target that was used to parametrize the neuromodulatory synaptic inputs to neocortical cells. The schema represents the main cell types in the neocortex. Abbreviations: PYR, pyramidal cell; INT, interneuron; SOM, somatostatin-positive cells; PV, pv-positive cells. **C)** Implementation of the synaptic model. Left: equations describing the Tsodyks-Markram model of synaptic short-term dynamics (above) and list of model parameters (below). Right: example of an in silico paired-recording experiment (excitatory, depressing connection) (above), and schematics illustrating the parameters that are modulated in order to mimic neuromodulatory effects. Gmax maximal conductance; Tau Decay PSC, decay time constant of the postsynaptic current; Use, utilization of synaptic efficacy; TauD, time constant of depression; TauF, time constant of facilitation; NRRP number of readily-released pool vesicles. **D)** Implementation of the volumetric transmission model for the three neuromodulatory systems. Graphs illustrating the distance-dependent change in Gmax. **E)** first, from above: schematics illustrating the spherical sampling portion of the algorithm developed to model volumetric transmission (VT). Second, partial validation of the VT model for ACh release (data replotted from Yamasaki et al, 2010). Third, partial validation of the VT model for DA release (data replotted from Courtney and Ford, 2014).

### Predicting ACh release kinetics

The dynamics of ACh release have not been elucidated yet, likely because they can be influenced by a variety of factors. ACh can be released tonically or phasically, via volume or synaptic transmission, and it can be subjected to fast clearing from the extracellular space because of the activity of catalytic enzymes such as cholinesterases (Coppola et al., 2016; Lysakowski et al., 1989). The literature reported values for the offset decay kinetics of ACh currents range across several orders of magnitude (Arroyo et al., 2012; Hay et al., 2016; Nelson & Mooney, 2016). To reconcile the conflicting reports, we simulated the cholinergic modulation of network activity as if it were 1) completely synaptic 2) wholly mediated by volumetric transmission and by varying the input type (phasic or tonic stimulation) and the kinetic parameters assigned to the connections to check the subsequent network effects. We designed a stimulus to mimic the phasic activation of cholinergic ascending fibers (a single-pulse, high frequency stimulus) and one to simulate a more tonic stimulation (a 20 Hz train of pulses) as was done experimentally by Hay and others (Hay et al., 2016). To select an appropriate volume of influence for the neuromodulatory volumetric signal we searched for studies reporting the spatial extent of neuromodulatory transients; often these are quite recent studies that leverage recently developed genetically-encoded-fluorescent sensors. To instantiate the ACh VT model, we assigned a value of 5 microns to the R_max_ parameter (see Methods and **Figure 4**), which was kept fixed for all simulations of volumetric cholinergic release, as reported by Borden and colleagues (Borden et al., 2020). The DTC values (**Table 6**) are important to determine the transmission timeline in both synaptic and volumetric release. While neuromodulatory ST is known to act on rapid timescales (i.e., the ms range), VT works with significantly longer transmission delays (hundreds of milliseconds) (Agnati et al., 2006; Arroyo et al., 2012). Nevertheless, synaptically mediated cholinergic currents with a slower decay have been observed as well (Hay et al., 2016). We therefore used some of the experimentally obtained DTC values and combined them with the other conditions in order to predict which set of parameters leads to a more substantial effect on network activity. We found that specific combinations of input type – transmission mode – and kinetic parameters are required to fully desynchronize the microcircuit (**Fig. 5**). Specifically, when ACh is released synaptically in our model, a 10 ms pulse can impact network activity only when the connections are assigned a slow enough decay kinetics (∼200 ms), while no effect can be observed when a faster kinetic is at play (∼ 5 ms). A 20 Hz train of pulses is even more impactful and leads to a prolonged desynchronization of network activity. However, the strongest desynchronization occurs when in our model ACh is released volumetrically (99% reduction; p-value = 9.09·10^−40^). In this case, if a train stimulation is coupled to even slower decay kinetic values (∼600 ms), activation of cholinergic fibers leads to a complete cessation of slow oscillatory phenomena, and network activity shifts to a much faster and entirely desynchronized regime as can be seen in **Fig. 6**. In our simulations cholinergic stimuli bring about a significant decrease of the delta band (1.5-3 Hz) power that is not due to a mere change in the mean firing frequencies of the neurons in the network that are receiving the stimuli. Therefore, all subsequent simulations implementing the VT model were assigned a DTC of 608.6 ± 109.7 ms (Hay et al., 2016) while for the simulations implementing the ST model, we used DTC = 241.2 ± 15.5 ms (Nelson & Mooney, 2016).

**Figure 5.**
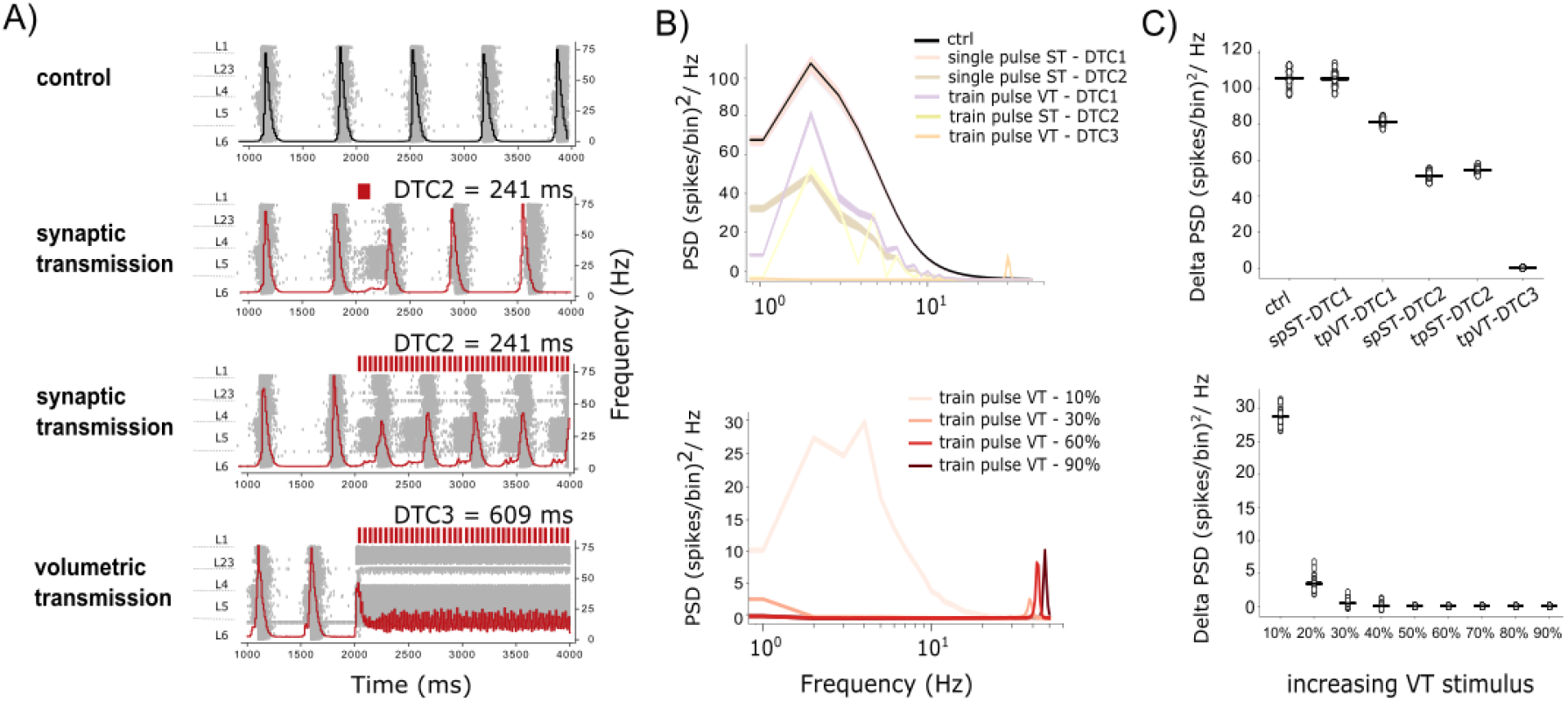
Network effects of volumetric or synaptic ACh release. Simulated network effects during the progressive activation of the virtual cholinergic projection systems. Timing of simulated optogenetic ACh release is shown as colored vertical bars on top of the plots. Simulation time is 4000 ms, and projection activation occurs at t = 2000 ms and stops at t = 4000 ms. **A)** Cholinergic effects; raster plots and superimposed frequency histograms. ST: synaptic transmission; VT: volumetric transmission; Ctrl: control condition. DTC: decay time constant of the PSC **B)** time-frequency representation plots. **C)** Graph showing only the delta (1.5-3 Hz) range power for every simulated condition. sp: single pulse; tp: train pulse.

**Figure 6.**
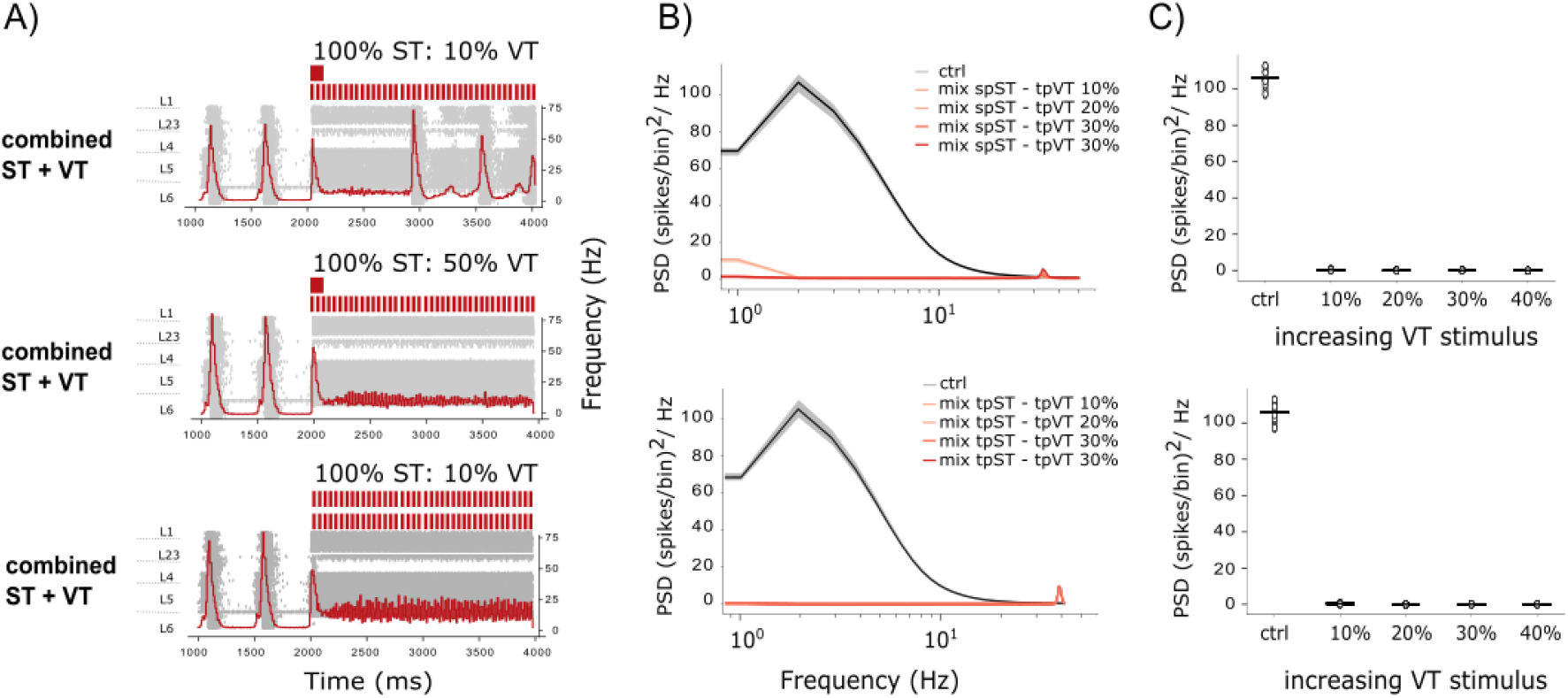
Network effects of combined volumetric and synaptic ACh release. Simulated network effects during the progressive activation of the virtual cholinergic projection systems. Timing of simulated optogenetic ACh release is shown as colored vertical bars on top of the plots. Simulation time is 4000 ms, and projection activation occurs at t = 2000 ms and stops at t = 4000 ms. **A)** Cholinergic effects; raster plots and superimposed frequency histograms. ST: synaptic transmission; VT: volumetric transmission; Ctrl: control condition. DTC: decay time constant of the PSC **B)** time-frequency representation plots. **C)** Graph showing only the delta (1.5-3 Hz) range power for every simulated condition. sp: single pulse; tp: train pulse.

### Relative contributions of synaptic and volumetric release

We showed that the VT implementation of cholinergic release dramatically desynchronized microcircuit activity while the ST implementation had a smaller effect on the reduction of slow oscillations. We therefore wondered what would happen if we activated the two systems simultaneously while manipulating the amount of input received. We performed an in-silico experiment where we only simulated cholinergic VT and activated increasing proportions of the afferent projections, thus simulating progressively higher levels of cholinergic release. For example, activating 10% of the projections in our model means that 28/283 source neurons are firing at 20 Hz. As can be seen in **Figure 6** we found that at least 50% of the VT projections have to be activated to completely desynchronize network activity. However, if we add synaptic release (100% of the fibers activated and firing at a 20 Hz frequency), lower levels of VT are sufficient to induce full network desynchronization (as low as 10% of VT), suggesting that the two modalities might work synergistically to evoke network states transitions. We kept testing this hypothesis, and performed another experiment where we activated progressively higher levels of VT while simultaneously simulating ST, this time by activating the ST projections with a high-frequency single pulse stimulation. Low levels of VT caused a complete desynchronization of microcircuit activity that was however shorter in duration (∼1 s) (**Figure 6**). The longer-lasting shift in network activity (comparable to results obtained with a tonic stimulation, ∼ 3 s) could be still rescued by recruiting higher levels of VT, once again suggesting that VT and ST might work together in a diversity of contexts.

### Simulating dopaminergic network effects

Dopamine is known to modulate arousal and promote wakefulness in living animals (Taylor et al., 2016), and to have effects on neocortical cell types (Bassant et al., 1990; Gonzalez-Islas & Hablitz, 2003; Gorelova et al., 2002). However, VTA inputs to sensory cortices have not been extensively described (as opposed to inputs to the prefrontal cortex), nor has their impact on network activity (unlike behavioral effects). For lack of better options, we used data about the effects on neocortical cell types elicited by bath application of dopaminergic agonists, mostly obtained in the prefrontal cortex, to parametrize the newly added dopaminergic synapses. As reported in **Table 4** and **Figure 4**, in our model, dopaminergic inputs depolarize L1 interneurons, all-layers proximal-targeting interneurons and pyramidal cells in L5 and L6. Pyramidal cells in L23 and L4 are instead inhibited by DA. We chose not to target the remaining cell-types because of lack of literature-reported effects. For a more detailed explanation of the parameters used for simulations, we redirect the reader to the Methods section of this paper and to **Table 6**. We implemented both the all-synaptic (ST) and the all-volumetric (VT) model of dopaminergic release, for which we used a Rmax = 2 μm. The DTCs values were constrained through literature-reported values [ST: 220 ± 41 ms (Condon et al., 2021); VT: 0.4 ± 0.1 s (Courtney & Ford, 2014). First, we simulated the all-synaptic activation of the dopaminergic projection system at increasingly higher stimulation rates (5, 10, 15, 20, 25, 30 Hz) to model progressively higher levels of dopamine release. As shown in **Figure 7** dopaminergic ST inputs significantly reduce the delta component of the power spectrum of network activity (∼26% reduction, p-value = 1.04·10^−21^). For all subsequent simulations of dopaminergic inputs, we chose a firing frequency of 20 Hz, again to be aligned with optogenetic stimulation experiments (Brunk et al., 2019; Taylor et al., 2016). Then we simulated an all-volumetric activation of dopaminergic projections and we found that VT transmission has a higher impact on network synchrony than ST: the delta power is reduced of 85% when 100% of the DA fibers are stimulated (p-value = 1.05·10^−26^) Furthermore, we did another experiment where we activated an increasingly higher proportion of inputs (10%, 20%, 30% up to 90% of the projections) thus simulating progressively increasing recruitment of neuromodulatory fibers. Dopaminergic inputs do not lead to a complete desynchronization of network activity, but the effect on slow oscillations seems to increase linearly with the number of fibers recruited. We also combined ST (100% fibers) with increasing amounts of VT (10, 20, 30 … 90% fibers) and found that when the two systems are activated simultaneously, low levels of VT are sufficient to induce complete desynchronisation in network activity (100% reduction); the effect remains stable at higher levels of VT.

**Figure 7.**
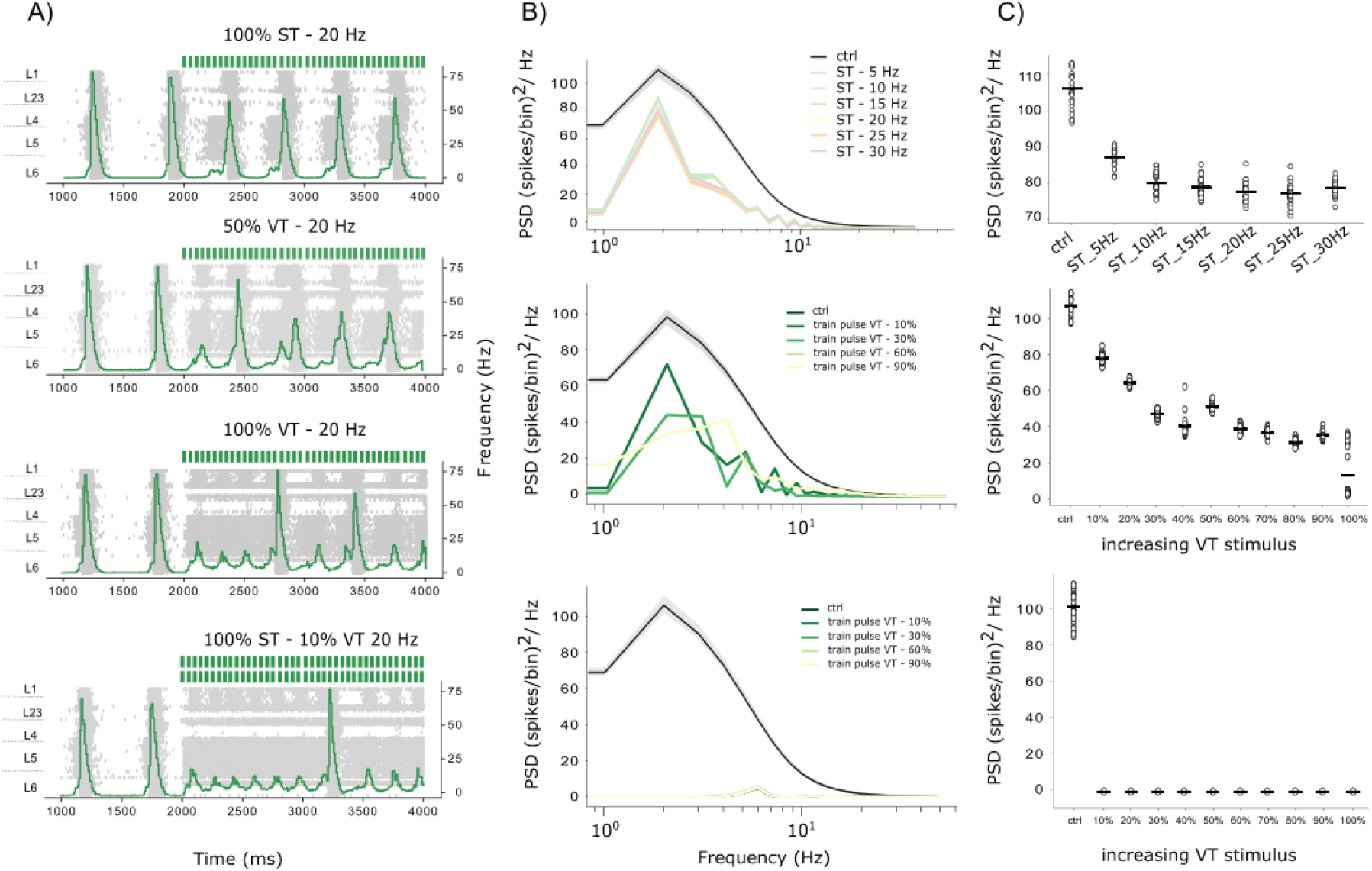
DA network effects. Simulated network effects during the progressive activation of the virtual dopaminergic projection systems. Timing of simulated optogenetic dopamine release is shown as colored vertical bars on top of the plots. Simulation time is 4000 ms, and projection activation occurs at t = 2000 ms and stops at t = 4000 ms. A) Dopaminergic effects; raster plots and superimposed frequency histograms. ST: synaptic transmission; VT: volumetric transmission; Ctrl: control condition. B) time-frequency representation plots. C) Graph showing only the delta (1.5-3 Hz) range power for every simulated condition.

### Simulating serotoninergic network effects

A substantial body of evidence suggests a causal relationship between 5-HT levels and cortical activity (Grandjean et al., 2019; Harris & Thiele, 2011; S.-H. Lee & Dan, 2012; Puig et al., 2010); optogenetic stimulation of the DR reduces low frequency power in the cortex thus promoting desynchronization. However, these results are mostly obtained in prefrontal areas, and the role of 5-HT projections to the SSCX is not clear to date. To test whether this holds true for the somatosensory cortex as well, we parameterized the serotoninergic connections as reported in experiments where 5-HT was bath-applied (see **Table 5**). As reported in **Figure 4 panel D**, serotoninergic inputs hyperpolarize lower layers (L4 and L5) pyramidal cells and L23 PV interneurons, but have excitatory effects on L23 VIP and SST interneurons, and (as for most neuromodulators) they depolarize all L1 interneurons. We chose not to target the remaining cell-types because of lack of literature-reported effects. For a more detailed explanation of the parameters used for simulations, we redirect the reader to the Methods section of this paper and to **Table 6**. We implemented both the all-synaptic (ST) and the all-volumetric (VT) model of 5-HT release, for which we used a Rmax = 3 μm. The DTCs values were constrained through literature-reported values (we assigned ST and VT the same value for lack of better data: 0.44 ± 0.3 s (Courtney & Ford, 2016). First, we simulated the all-synaptic activation of 5-HT fibers with progressively increasing stimulation frequencies (5, 10, 15, 20, 25, 30 Hz) to check the effect on slow oscillations. We found that 5-HT inputs reduce the delta component of network oscillations (∼80% reduction, p-value = 1.6·10^−37^) and shift the power spectrum towards higher-frequency components; while the reduction of the 2 Hz peak is drastic, an increase in oscillation frequency is also evident, as can be seen in the raster plots in **Figure 8**. For all subsequent simulations of 5-HT release we used a train of pulses at 20 Hz (again, to fall in line with optogenetics experiments such as the study performed by (Lottem et al., 2016). The effect on 2 Hz oscillations is even larger when we simulate an all-volumetric implementation of 5-HT release (reduction of 99% p-value = 2.6·10^−40^), and it increases with the number of fibers stimulated (again, we simulated the activation of 10%, 20% … up to 100% of the fibers). The VT model of serotoninergic inputs also predicts an increase in higher-frequency oscillations (the theta range) and brings about a significant desynchronization of network activity. We also tried coupling the ST and VT models (where we activated 100% of ST inputs, and increasing percentages of VT release) and found that when the two release modes are combined, low levels (10%) of VT are sufficient to induce a complete desynchronization of network activity (100% reduction).

**Figure 8.**
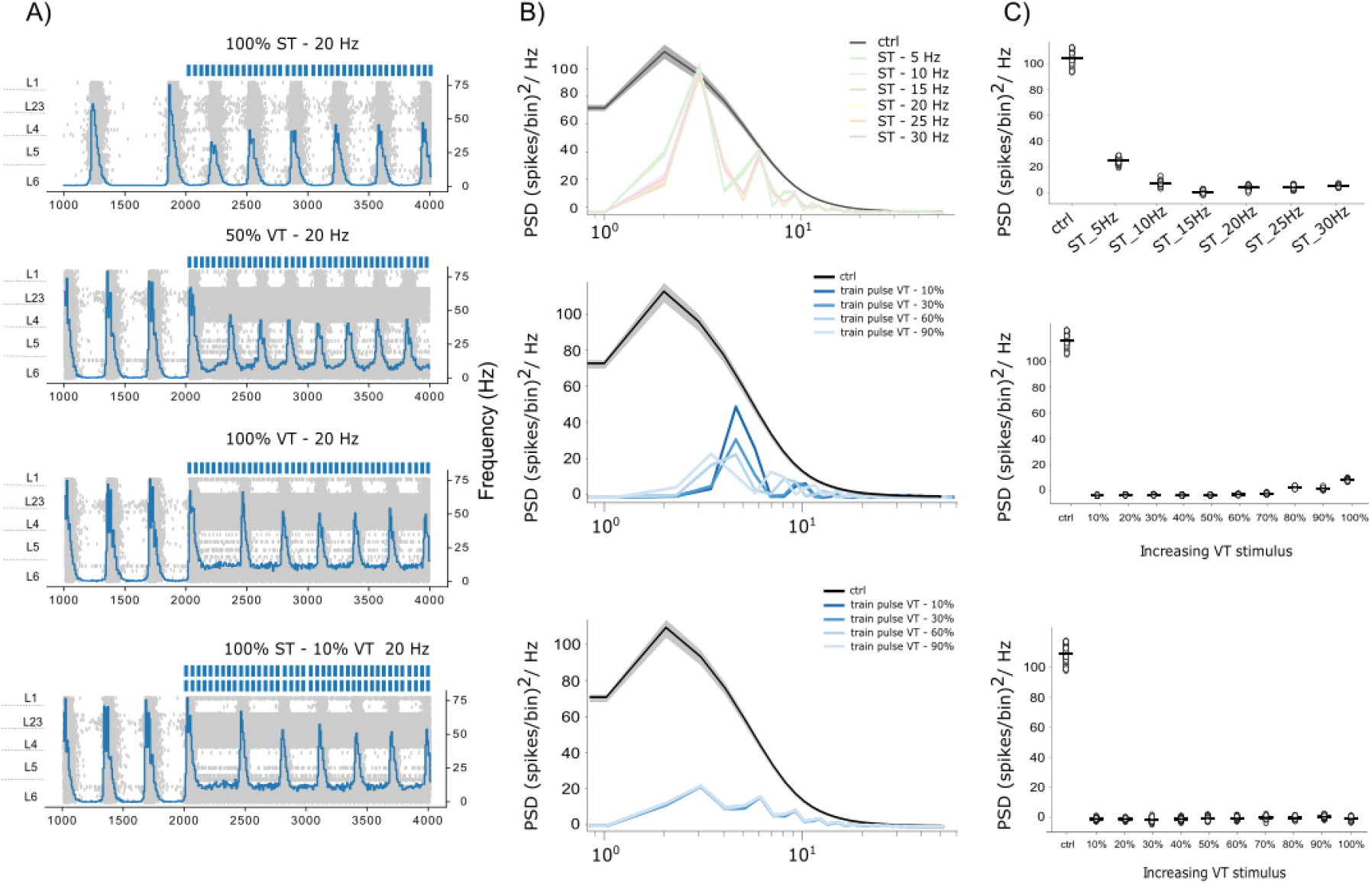
5-HT network effects. Simulated network effects during the progressive activation of the virtual serotoninergic projection systems. Timing of simulated optogenetic serotonin release is shown as colored vertical bars on top of the plots. Simulation time is 4000 ms, and projection activation occurs at t = 2000 ms and stops at t = 4000 ms. **A)** 5-HT effects; raster plots and superimposed frequency histograms. ST: synaptic transmission; VT: volumetric transmission; Ctrl: control condition. **B)** time-frequency representation plots. **C)** Graph showing only the delta (1.5-3 Hz) power range for every simulated condition.

## Discussion

### Anatomy of neuromodulatory systems

In the present study we have estimated densities and laminar distribution patterns of neuromodulatory systems in the rat P14 HLS1 using immunostaining and stereological techniques. The data are compared to previous data generated in our laboratory (**Table 2**) using FIB/SEM and Espina software that allow the identification and quantification of virtually all cortical synapses in long series of images that represent a 3D sample of cortical neuropil (Merchan-Pérez, 2009; Santuy et al., 2018). Previous studies in different cortical regions and species have examined the developmental maturation of the rat cortical innervation by the networks of cholinergic, catecholaminergic and serotonergic fibers (Descarries & Mechawar, 2000; Dori et al., 1996; Kalsbeek et al., 1988; Latsari et al., 2002; Lidov & Molliver, 1982; Mechawar et al., 2002; Mechawar & Descarries, 2001; Verney et al., 1984). These studies suggested that the pronounced effects that these regulatory systems exert on the morphology and physiology of maturing cortical neurons are both through synaptic and non-synaptic connections. They reported regional specificity throughout the neocortex and progressive changes over postnatal development in fiber length, branching patterns, number of varicosities and percentage of varicosities that form synaptic contacts. To get a deeper insight in the organization of the cortical neuropil in the different cortical layers of the HLS1 of two-week old rats, in the present study we have also estimated the fiber length per cortical volume and the density of varicosities of catecholaminergic, serotonergic and cholinergic systems, using immunocytochemical and stereological techniques. Our combined experimental and modeling approach demonstrates that the cholinergic projection system prevails over other neuromodulatory systems in the cerebral cortex and contacts by far the largest number of postsynaptic targets in each neocortical layer. The second most prevalent type of neuromodulatory innervation is the TH-positive fiber system. Dopaminergic fibers are generally thought to be present mostly in the frontal areas (Jacob & Nienborg, 2018) while only sparsely innervating the sensory cortices (Aransay et al., 2015). Our findings confirm previous results concerning the presence of TH staining in the developing somatosensory cortex (Descarries et al., 1987). Anatomical studies in different rodent species have identified substantial serotonergic projections from the raphe nuclei to early sensory areas including the somatosensory cortex (Jacob & Nienborg, 2018). 5-HT fiber density transiently increases in the neonatal rodent neocortex (D’Amato et al., 1987), and becomes more uniformly distributed after the third postnatal week. Here, we confirm that serotoninergic fibers innervate the developing rodent neocortex and are more abundantly present in superficial rather than deep layers. This preferential organization of serotoninergic fibers and varicosities could reflect the fact that a large, developmentally distinct category of inhibitory interneurons that also expresses the 5HT3aR is concentrated in supragranular layers. This population is heterogeneous and includes all of the VIP expressing neurons, as well as an equally numerous subgroups of neurons that do not express VIP and includes neurogliaform cells (Rudy et al., 2011). The preferential innervation of L1 seems to be a key feature of neuromodulatory projections to the somatosensory cortex; given the relative scarcity of cell bodies in layer 1 it is reasonable to hypothesize that they would be contacted by a large number of fibers, in order to maximize the probability of receptor activation. To gain a better insight of the organization of neuromodulatory systems we also developed a structural model of cholinergic, dopaminergic and serotoninergic innervation of the sensory cortex. Our goal here was to generate predictions of quantities that are hardly measurable in real-life settings such as the number of neurons contacted by each neuromodulatory axon, the number of synapses established by each fiber and the proportions of innervated cell-types.

**Table 2.** Correlations between varicosities and neuronal densities. Correlations between the mean values in each cortical layer of vGlut1-ir, vGlut2-ir and vGAT-ir axon terminals, 5HT-ir, TH-ir and ChAT-ir fiber and varicosities density (var), neuronal density (**ρ)** (NeuN-ir, taken from (Markram et al., 2015)) and density of symmetric (SS) and asymmetric (AS) synapses (modified from (Merchán-Pérez et al., 2014)). Numbers indicate R^2^ values (Pearson test); p-values are shown between brackets.

| vGlut<br>1 | vGlut<br>2 | vGAT | ρSer | ρTH | ρCHAT | Var<br>Ser | Var<br>TH | Var<br>CHAT | ρNeu | SS | AS |  |
| --- | --- | --- | --- | --- | --- | --- | --- | --- | --- | --- | --- | --- |
|  | 0.81<br>(0.01) | 0.77<br>(0.02) | 0.53<br>(0.09) | 0.58<br>(0.07) | 0.52<br>(0.1) | 0.62<br>(0.06) | 0.77<br>(0.02) | 0.96<br>(.0004) | 0.39<br>(0.18) | 0.22<br>(0.34) | 0.28<br>(0.27) | vGlut1 |
|  |  | 0.8<br>(0.01) | 0.42<br>(0.16) | 0.75<br>(0.02) | 0.35<br>(0.21) | 0.45<br>(0.14) | 0.79<br>(0.01) | 0.68<br>(0.04) | 0.29<br>(0.27) | 0.46<br>(0.13) | 0.43<br>(0.42) | vGlut2 |
|  |  |  | 0.42<br>(0.16) | 0.95<br>(0.0009) | 0.17<br>(0.41) | 0.43<br>(0.15) | 0.86<br>(0.007) | 0.65<br>(0.05) | 0.13<br>(0.48) | 0.66<br>(0.05) | 0.31<br>(0.24) | vGAT |
|  |  |  |  | 0.35<br>(0.21) | 0.58<br>(0.07) | 0.97<br>(.0003) | 0.55<br>(0.08) | 0.47<br>(0.13) | 0.18<br>(0.39) | 0.04<br>(0.68) | 0.14<br>(0.46) | ρSer |
|  |  |  |  |  | 0.07<br>(0.59) | 0.33<br>(0.23) | 0.79<br>(0.01) | 0.44<br>(0.14) | 0.05<br>(0.65) | 0.8<br>(0.01) | 0.31<br>(0.25) | ρTH |
|  |  |  |  |  |  | 0.72<br>(0.72) | 0.42<br>(0.42) | 0.58<br>(0.58) | 0.75<br>(0.74) | 0.01<br>(0.02) | 0.00<br>(0.008) | ρCHAT |
|  |  |  |  |  |  |  | 0.6<br>(0.06) | 0.58<br>(0.07) | 0.3<br>(0.25) | 0.02<br>(0.75) | 0.09<br>(0.54) | VarSer |
|  |  |  |  |  |  |  |  | 0.7<br>(0.03) | 0.4<br>(0.17) | 0.4<br>(0.18) | 0.1<br>(0.53) | VarTH |
|  |  |  |  |  |  |  |  |  | 0.49<br>(0.12) | 0.12<br>(0.49) | 0.17<br>(0.41) | VarCHAT |
|  |  |  |  |  |  |  |  |  |  | 0.0<br>(0.89) | 0.02<br>(0.75) | ρNeu |
|  |  |  |  |  |  |  |  |  |  |  | 0.31<br>(0.24) | SS |
|  |  |  |  |  |  |  |  |  |  |  |  | AS |

### Physiology of neuromodulatory systems

Neuromodulators are traditionally thought to act with low spatial precision throughout the cortex, but recent lines of evidence suggest that they also exhibit fast modes of signaling (Kalmbach et al., 2012; Obermayer et al., 2018; Poorthuis et al., 2013) and that neuromodulatory axons establish specialized synaptic contacts (Nelson & Mooney, 2016; Takács et al., 2013; Turrini et al., 2001). In rodent neocortex reports of the percentage of varicosities that establish synaptic contacts are conflicting and values range from 14% to 66% (Colangelo et al., 2019). Some authors argue that cholinergic synapses might be difficult to find with traditional methods, and propose novel ways in which they could be identified (Nelson & Mooney, 2016; Takács et al., 2013; Turrini et al., 2001); Tak(Nelson & Mooney, 2016; Takács et al., 2013; Turrini et al., 2001)cs and colleagues for instance observed that cholinergic contact sites in rodent neocortex were strongly labeled with neuroligin-2 and that they did not resemble typical synapses, suggesting that cholinergic fibers establish more synaptic connections than it was previously recognized. More in general, a body of anatomical evidence suggests that cholinergic synapses exist, with apposition of pre and postsynaptic sites, although they might not account for all ACh release sites (Disney & Higley, 2020; Turrini et al., 2001). Evidence in favor of cholinergic volumetric transmission, such as the presence of extrasynaptic receptors and slowly decaying current kinetics (Descarries & Mechawar, 2000; Hay et al., 2016; Yamasaki et al., 2010) has also been reported. According to Hay and others (Hay et al., 2016), both synaptic and diffuse cholinergic transmission occur in the neocortex depending on the regime of BF neuronal activity. Thus, even though fast point-to-point modulation of synaptic transmission has been repeatedly demonstrated in the rodent neocortex (Kalmbach et al., 2012), the possibility that neuromodulatory systems exert their action via volume transmission as well cannot be excluded. Incidentally, whilst *en passant* axonal boutons of cholinergic neurons of subcortical provenance can mediate volume transmission broadly in the cortex, acetylcholinesterase (AChE) restricts the diffusion of ACh by enzymatic hydrolysis after its release (Sarter et al., 2009). Thus, slow and diffuse cholinergic transmission in the cortex is likely to be spatially and temporally constrained by the catalytic power of AChE, other than by the localization and density of cholinergic receptors. It is reasonable to hypothesize that neuromodulatory activity likely reflects a mixture of spatiotemporal dynamics (Muñoz & Rudy, 2014); however, the relative contributions of the two transmission modalities remain unclear. In this study, we implemented a model of neuromodulatory release that accounts for volumetric and synaptic transmission to investigate the dynamics of cholinergic release in sensory cortices. Additionally, we extend our framework to include other neuromodulators, such as DA and 5-HT, that have not been so extensively studied in the somatosensory cortex of the rat. We show that cholinergic projections modulate the activity of the simulated microcircuit by decreasing slow oscillations (delta oscillations) and by desynchronizing network activity. In particular we show that high levels of volumetric transmission are sufficient to fully and persistently desynchronize the microcircuit, but they are not critically required: when VT is coupled to ST, we observe a synergic effect and lower levels of VT suffice to achieve the same network effect. Moreover, we show that conflicting reports about the magnitude of the DTC in cholinergic induced PSCs can be reconciled via *in silico* experiments performed within our framework, and so can the cell-type specific effects of cholinergic release. No amount of validation can prove a model right, but within its domain of validity our model allowed us to put to the test the underlying hypothesis (i.e. the data that we used to constrain the model). The literature about the cell-type specific cholinergic modulation of membrane properties is quite conflicting. For example, there is no clear agreement on the effects exerted by ACh on pyramidal neurons, nor on basket cells (PV-FS interneurons). Some papers reported inhibitory effects, others excitatory or biphasic, or (sometimes paradoxically) a lack of effect (Eggermann & Feldmeyer, 2009; Gulledge & Stuart, 2005; Hedrick & Waters, 2015; Joshi et al., 2016; Obermayer et al., 2018; Xiang et al., 1998). The same is true for measurements of transmission features such as the DTC, or the firing frequency of cholinergic cells of subcortical provenance. Because our model replicates well-established emergent phenomena such as the desynchronizing effect of cholinergic release reported in the literature (Alitto & Dan, 2013; Pinto et al., 2013), we can be confident that it captures at least some essential properties of the system being modeled. Less is known about the activation of the DA and 5-HT modulatory systems and their effects on sensory microcircuits. Most of the data is recorded in prefrontal regions or in subcortical modulatory regions such as the striatum and the DR (Courtney & Ford, 2016; Gonzalez-Islas & Hablitz, 2003; Puig et al., 2010), so it’s harder to validate the results of our model. However, the presence of both DA and 5-HT varicosities and receptors has been reported in the rodent sensory cortex (Borden et al., 2020; D’Amato et al., 1987; Descarries et al., 1987). We decided to extend the assessment of the influence of these neuromodulatory systems in otherwise previously unexplored conditions, i.e., in somatosensory regions. We simulated the effects of endogenous DA release and we found that synaptic transmission alone induces a significant decrease in the delta power (30%) of microcircuit oscillatory activity; VT however is much more impactful in that it has an even stronger effect of slow oscillations. Combining dopaminergic VT and ST leads to complete network desynchronization. It is interesting to notice here that the desynchronization induced by DA is not as long-lasting as seen in the ACh case even though the train stimulation is equivalent. 5-HT volumetric and synaptic inputs also drastically reduced the delta component of network oscillations and cooperated to bring about complete network desynchronization. Additionally, serotoninergic release induced a shift in the power spectrum towards higher-frequency components (5-8 Hz).

### Modeling assumptions

When modeling, one is necessarily simplifying from ground truth. These simplifications are based on assumptions, where as long as the assumption holds true, the corresponding simplification does not limit the validity of the model. Consequently, in order to assess the validity of a model it is paramount to be aware of the assumptions it is based on. In the next paragraph we will describe three main types of assumptions that our model is based upon. The first type is the “inherited assumption”, that occurs whenever a model is built on top of another existing model. The new model inherits all relevant assumptions of the base model. For example, our algorithm to reconstruct neuromodulatory innervation inherits the assumptions relating to neuronal composition, placements and morphology from the underlying neocortical column model (Markram et al., 2015). The second type is the data/structuring assumption: that is, an assumption that acts on how the model is parameterized, i.e., how data is turned into parameters. In this modeling study, after placing additional synapses in our NCX model to match the distribution profile observed experimentally, we assigned synaptic features based on literature reports gathered in **Tables 3-4-5**. We developed an algorithm that assigns excitatory or inhibitory synaptic parameters based on the postsynaptic morphological type (m-type) contacted by our virtual projection systems. We assumed that synapses of subcortical provenance will display forms of short-term dynamics similar to the well-known dynamics of neocortical synapses (Markram et al., 2015) and that they would be governed by similar principles. Moreover, we assume that organizational principles extracted for specific cell-types hold true in more general cases and we apply them to broader target classes. For example, if we find that ACh depolarizes L5 pyramidal cells we extend this observation to a broader category, i.e., L5 excitatory cells. Additionally, we compute the number of fibers for each neuromodulatory projection system based on rough estimations of the number of presynaptic cells in the underlying subcortical neuromodulatory region. This assumption has limited validity in many ways. 1) We are assuming that NM projections originate from a specific nucleus in the subcortical modulatory region (in particular: the NBM for ACh, the DR for 5-HT and the VTA for DA) and we are disregarding the possibility that some proportion of the connections can arise from other less relevant or less studied nuclei. While it is true that these nuclei are the main sources of neuromodulatory inputs, the possibility that other nuclei are also involved cannot be excluded. 2) We assume that the innervation of the S1 region is homogeneous in all its subregions, while there may be differences in the innervation of specific areas. 3) We report modeling assumptions, that is all the assumptions at the very core of the model itself. For instance, we assume that the slow oscillatory activity of our simulated microcircuit (yielded by a specific set of parameters) can mimic the inactivated/synchronized brain state typical of SWS states (see Methods section – Microcircuit). The third type of modeling assumptions revolve around the VT implementation. Information such as the concentration of the neurotransmitter in the extracellular matrix, and the dynamics of the release in terms of diffusion kinetics is central to understanding the spatiotemporal constraints of VT. However, neither the effective concentration of neuromodulators in the extracellular space, nor the rates of their diffusion/degradation are known and an accurate knowledge of the spatiotemporal limits of diffuse neuromodulator release is still lacking (Coppola et al., 2016), mostly because of technological limitations. In our model, we assume that the extracellular matrix has no influence on volumetric transmission, and we do not specifically model the action of catalytic enzymes like cholinesterases. Additionally, we sample a spherical volume around the VT release site and assume transmission isotropy. A list of assumptions can be complete only when the following conditions are met: the infinite set of all conceivable models must be considered, and then each assumption, combined with ground truth data, rules out a part of that set. Then, the proposed model is the only one remaining that is consistent with the ground truth data. The theoretical goal of having a complete list of assumptions is therefore unachievable, for the simple reason that the space of all conceivable models is most likely impossible to describe, but it provides a useful thought framework.

**Table 3.** Summarized literature reported cell-type specific effects of cholinergic release. Inclusion criteria privilege studies performed a) by means of optogenetic stimulation of subcortical nuclei as a method to evoke neuromodulator release; b) in the somatosensory areas of the rodent neocortex c) in developing brains. When this is not applicable (i.e., data is missing) data was taken from studies using electrical stimulation of subcortical nuclei or bath-application of neuromodulators as methods to evoke neuromodulator release.

| ChAT | Pyramidal cells | Interneurons | Reference |
| --- | --- | --- | --- |
| **Layer 1** |  | EPSP (and spiking activity) | (Arroyo et al., 2012) (p20-40 mice SSC) optogenetics |
|  |  | EPSC | (Hay et al., 2016) (p25 mice);<br>(Bennett et al., 2012) (adult mice);<br>optogenetics |
| **Layer 2/3** | IPSC barrage | EPSP (BPCs) | (Arroyo et al., 2012) (p20-40 mice SSC) optogenetics |
|  |  | SOM increase FF; PV reduced FF | (Chen et al., 2015) (mouse v1) optogenetics |
| **Layer 4** | IPSP |  | (Meir et al., 2018) (adult mice SSC) optogenetics |
| **Layer 5** | EPSP or IPSP |  | (Hedrick & Waters, 2015) (mouse V1) optogenetics |
|  | EPSP or IPSP |  | (Joshi et al., 2016) (p24 mouse A1) optogenetics |
| **Layer 6** | EPSC |  | (Hay et al., 2016) (p25 mice);<br>(Verhoog et al., 2016) (mouse PFC) optogenetics |

**Table 4.** Summarized literature reported cell-type specific effects of dopaminergic release. Inclusion criteria privilege studies performed a) by means of optogenetic stimulation of subcortical nuclei as a method to evoke neuromodulator release; b) in the somatosensory areas of the rodent neocortex c) in developing brains. When this is not applicable (i.e., data is missing) data was taken from studies using electrical stimulation of subcortical nuclei or bath-application of neuromodulators as methods to evoke neuromodulator release.

| TH | Pyramidal cells | Interneurons | Reference |
| --- | --- | --- | --- |
| **Layer 1** | | depolarization | (Zhou & Hablitz, 1999) (adult rat PFC) Bath application of DA (30–100 μM) |
| **Layer 2/3** | Hyperpolarization; depolarization | increase excitability of PV-FS | (Zhong et al., 2020) (3w rats PFC) in low Ca; (Zhou & Hablitz, 1999) (adult rat PFC) Bath application of DA (30–100 μM) |
| | | increase excitability of FS and depolarization | (Gorelova et al., 2002) (rat PFC) 40 μM DA |
| **Layer 4** | Hyperpolarization; depolarization | increase excitability of FS and depolarization | (Gorelova et al., 2002) (rat PFC) 40 μM DA; (Zhou & Hablitz, 1999) (adult rat PFC) |
| **Layer 5** | increase excitability via D1r (or sometimes decrease via D2r) | increase excitability of PV-FS | (Radnikow & Feldmeyer, 2018) (rodents PFC); (Zhong et al., 2020) (3w rats PFC); (Gao & Goldman-Rakic, 2003) (ferret PFC) |
| | | increase excitability of FS and depolarization; | (Gorelova et al., 2002) (40 μM DA, rat PFC); (Zhou & Hablitz, 1999) (adult rat PFC) |
| **Layer 6** | increase excitability |  | (Radnikow & Feldmeyer, 2018) (rodents PFC); (Zhou & Hablitz, 1999) (adult rat PFC) |

**Table 5.**
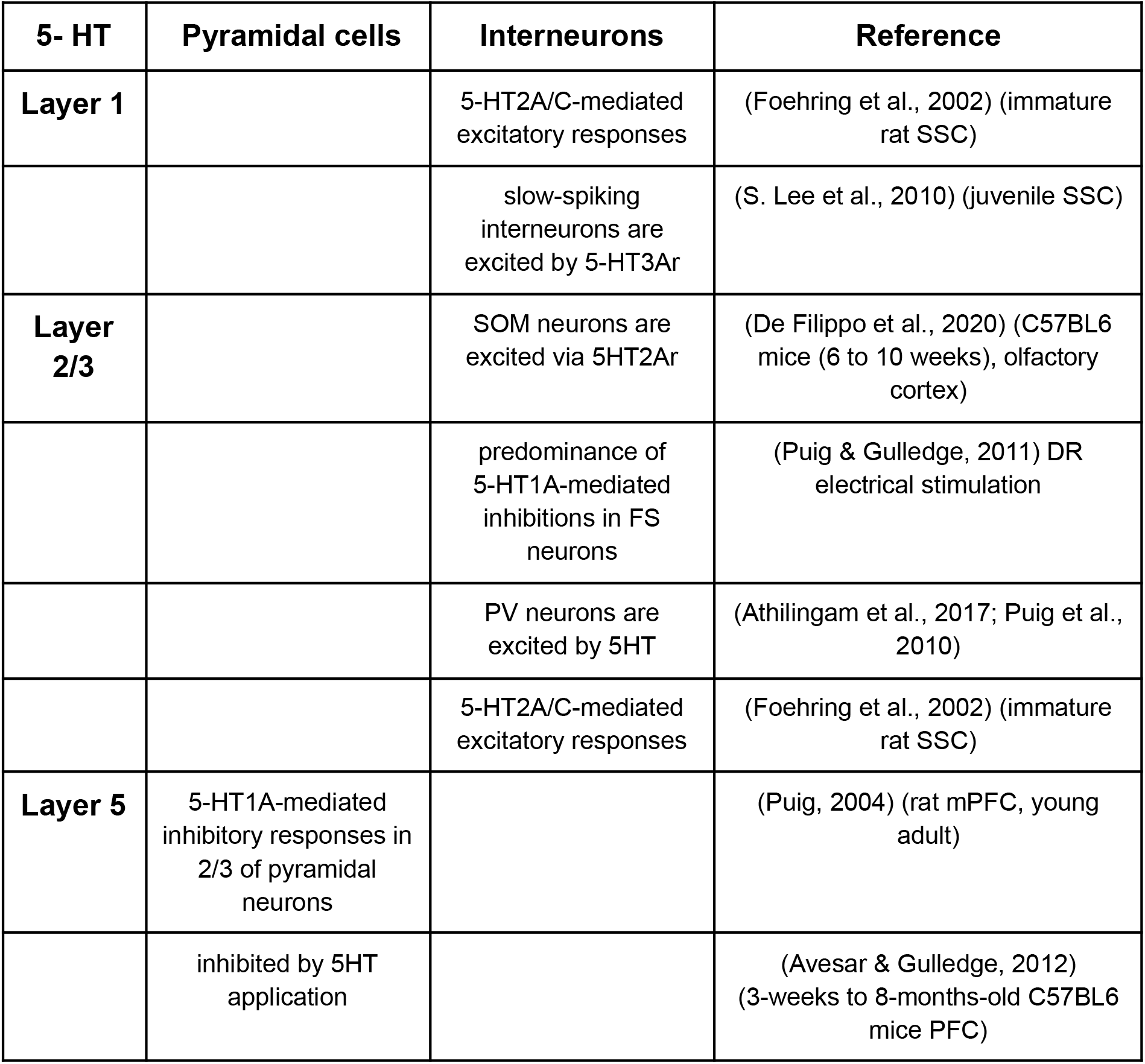
Summarized literature reported cell-type specific effects of serotoninergic release. Inclusion criteria privilege studies performed a) by means of optogenetic stimulation of subcortical nuclei as a method to evoke neuromodulator release; b) in the somatosensory areas of the rodent neocortex c) in developing brains. When this is not applicable (i.e., data is missing) data was taken from studies using electrical stimulation of subcortical nuclei or bath-application of neuromodulators as methods to evoke neuromodulator release.

**Table 6.**
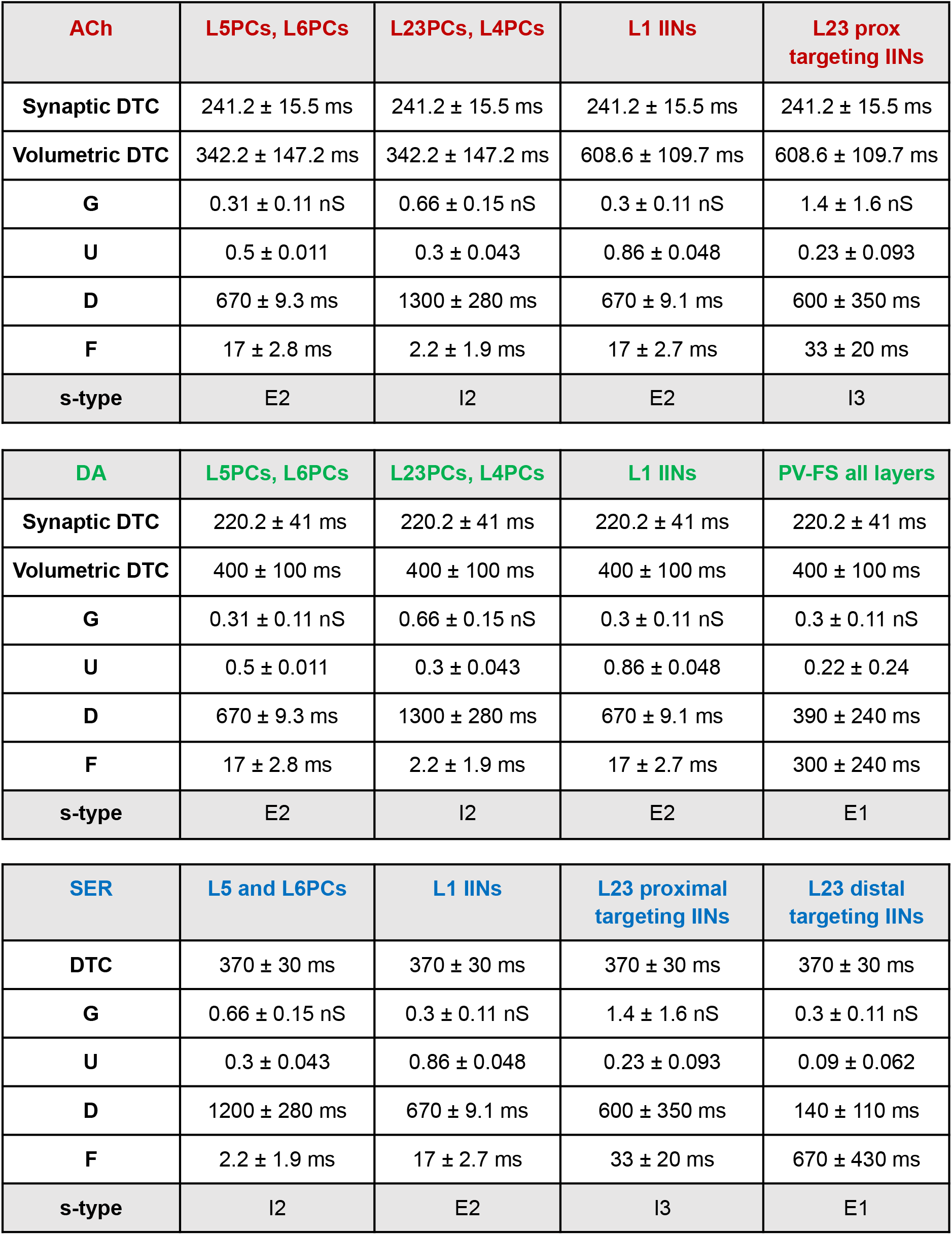
Neuromodulatory projections parameters. DTC: decay time constant of the PSC; G: synaptic conductance; U: time constant of synaptic depression; F: time constant of synaptic facilitation: s-type: synaptic type according to (Markram et al., 2015).

## Conclusions and future directions

Our goal here was to integrate sparse experimental datasets and provide a way to generate predictions about the influences of three distinct neuromodulatory systems in the somatosensory cortex. Through our modeling efforts we sought to provide a framework to investigate the long-standing issue of the relative importance of volumetric vs point-to-point synaptic transmission. Our results show that the two modalities have cooperative effects, and we propose that one key feature of network transitions between synchrony and asynchrony regimes is the co-occurrence of volumetric and synaptic neuromodulatory transmission. Predictably, our model has limitations, several of which we have already listed in the assumptions section. To mention a few others, we do not account for the action of other neuromodulators in the cortex (Disney, 2021), which are likely to influence the context in which ACh, DA and 5-HT work, nor we take into consideration the effects that these three neuromodulators have on each other and on their own activity. ACh is known to be involved in feedback regulation of cholinergic release via presynaptic axonal autoceptors for instance, and neuromodulators can be co-released with other peptides or transmitters (Granger et al., 2017; Saunders et al., 2015). The framework we built to model neuromodulatory transmission can be updated as we gather new data and move forward in our understanding of neuromodulatory systems.

## Supplementary Figures

**Supplementary Figure S1.**
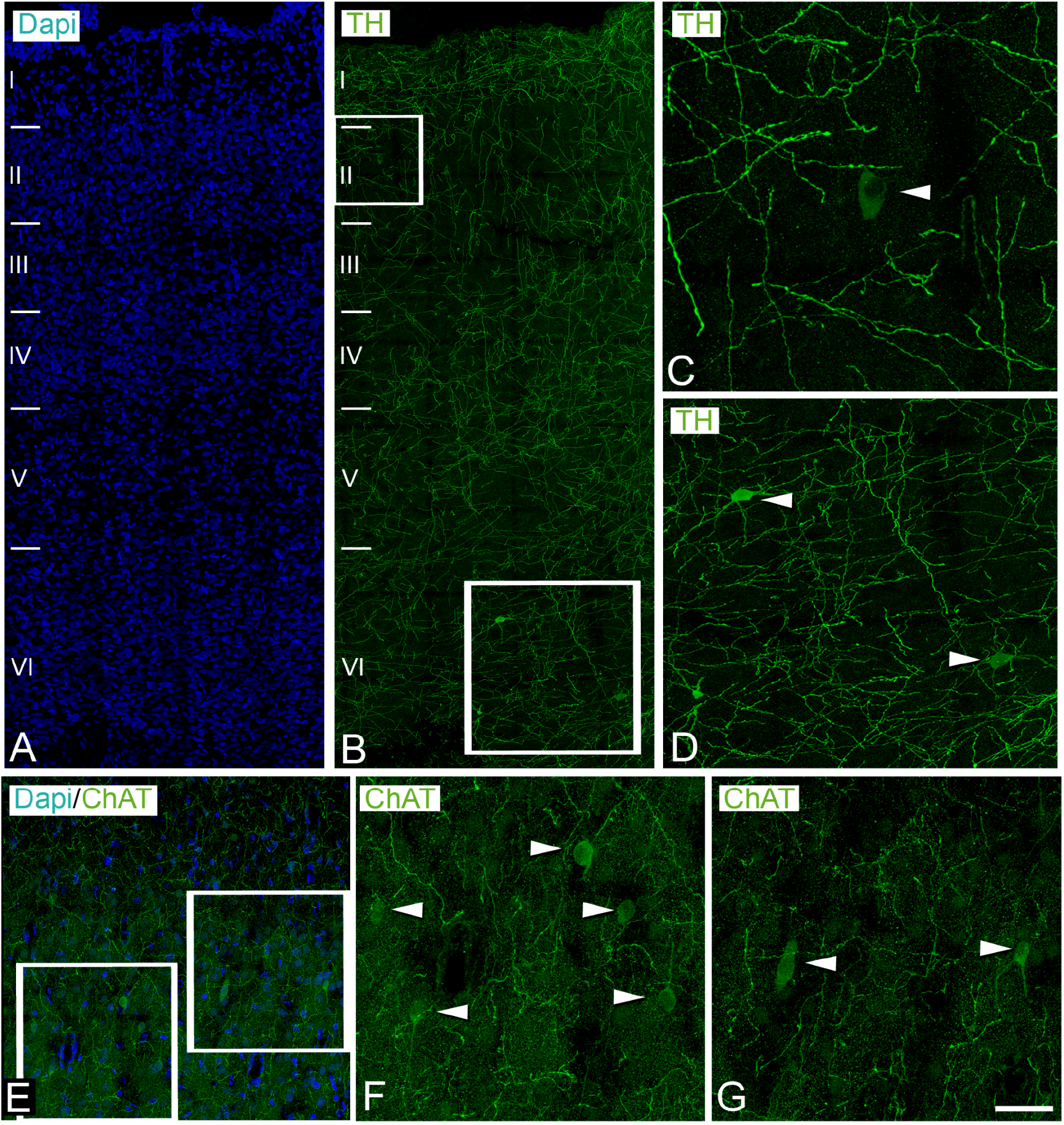
ChAT and TH intracortical cell bodies. **A)** confocal stack projection image, corresponding to a cortical thickness of 14 μm, showing DAPI staining in the different layers of the P14 rat hindlimb somatosensory cortex. **ti** same as in A but showing the distribution of TH-immunoreactive fibers (green). Squared zones are shown at higher magnification in panel **C)** and panel **D)** where arrowheads point to cell bodies. **E)** Same as in B) but for ChAT immunoreactive fibers. Squared zones are shown at higher magnification in panel **F)** and panel **G)** where arrowheads point to cell bodies. Scale bar spans 15 μm.

